# Supporting evidence for DNA shearing as a mechanism for origin unwinding in eukaryotes

**DOI:** 10.1101/739557

**Authors:** Lance D. Langston, Roxana Georgescu, Mike E. O’Donnell

## Abstract

Our earlier study demonstrated that head-to-head CMGs that encircle duplex DNA and track inward at origins, melt double-strand (ds) DNA while encircling the duplex by pulling on opposite strands and shearing DNA apart (Langston and O’Donnell (2019) eLife <u>9</u>, e46515). We show here that increasing the methylphosphonate neutral DNA from 10 nucleotides in the previous report, to 20 nucleotides, reveals that CMG encircling duplex DNA only interacts with the tracking strand compared to the non-tracking strand. This significantly enhances support that CMG tracks on duplex DNA by binding only one strand. Furthermore, EMSA assays using AMPPNP to load CMG onto DNA shows a stoichiometry of only 2 CMGs on an origin mimic DNA, containing a 150 bp duplex with two 3 prime single-strand (ss) tails, one on each end, enabling assay of dsDNA unwinding by a shearing force produced by only two head-to-head CMGs. The use of non-hydrolysable AMPPNP enabled a preincubation for CMG binding the two 3 prime tailed origin mimic DNA, and gave robust unwinding of dsDNA by head-to-head CMG-Mcm10’s. With this precedent, it is possible to envision that the cell may utilize opposing dsDNA motors to unwind DNA for other types of DNA transactions besides origin unwinding.

## INTRODUCTION

All cells utilize factors that assemble two helicases onto DNA for bidirectional replication forks. In *Escherichia coli* the DnaA origin binding protein is primarily responsible for the initial opening of double-strand (ds) DNA, upon which two DnaB hexameric helicase rings are assembled onto opposite strands of the single-stranded (ss) DNA bubble (reviewed in (Bleichert et al., 2017; O’Donnell et al., 2013)). Unlike bacteria, studies in *Saccharomyces cerevisiae* reveal that eukaryotic helicases are assembled on dsDNA. Helicases are assembled in two different stages of the cell cycle ((reviewed in (Bell and Labib, 2016; Bleichert et al., 2017; Parker et al, 2017)). In G1, origins are “licensed” by the Origin Recognition Complex (ORC), Cdc6, and Cdt1 that assemble two inactive Mcm2-7 motor rings around dsDNA (Evrin et. al., 2009; Reemus et al., 2009). In S-phase, two cell cycle kinases (DDK, CDK), Sld2, Sld3, Sld7, Dbp11, and Pol *ε* assemble Cdc45 and the GINS tetramer onto Mcm2-7 to form a complex of each of Cdc45, Mcm2-7, and GINS tetramer. Isolation and characterization of the 11-subunit CMG (Cdc45, Mcm2-7, GINS) complex demonstrated that CMG is the active form of the cellular helicase (Ilves et al., 2010; Moyer et al., 2006). The separation of active helicase into two cell cycle phases ensures that replication of a eukaryotic genome, containing numerous origins, occurs once and only once per cell cycle.

All replicative ring-shaped helicases studied thus far function by encircling ssDNA, including CMG (reviewed in (Lyubimov et al., 2011; O’Donnell and Li, 2018)). Thus, it has been a conundrum as to how CMGs melt origin DNA while encircling dsDNA. A cryoEM study revealed that the CMGs are directed to track inward, toward one another at the origin, and thus block one another (Georgescu et al., 2017). While this inward directed orientation of CMGs seems counterintuitive, it has been confirmed by three other studies thus far (Douglas et al., 2018; Goswami et al., 2018; Meagher et al., 2019). A recent study identified that the inward orientation of head-to-head CMGs on dsDNA held the key to how origin dsDNA is melted, and would not occur with CMG having an opposite orientation (Langston and O’Donnell, 2019). The mechanism involved CMGs binding one strand of dsDNA more tightly than the other, and therefore two opposed CMGs can pull on opposite strands shearing the duplex apart, providing the 65 pN force needed to melt dsDNA (King et al., 2016; van Mameren et al., 2009). Forces of this scale have precedent in oligomers that encircle DNA, the FtsK chromosome partitioning motor of E. coli (Pease et al., 2005) and the phi29 DNA packaging motor (Smith et al., 2001). The current report builds upon these observations by determining the stoichiometry of CMG during shearing, developing a robust melting/shearing reaction, and providing clear evidence of the strand asymmetry of CMG binding only one strand of DNA while encircling the duplex.

## RESULTS

### CMG binds the 3’-5’ strand while encircling dsDNA

To determine the extent to which one strand(s) of dsDNA is needed more than the other strand for CMG tracking with force while encircling dsDNA, we used a T-DNA containing a 3’ oligo dT tail for CMG loading followed by a flush duplex that is demonstrated to enable CMG to track over the ssDNA and onto the dsDNA (Kang et al., 2012; Langston and O’Donnell, 2017a). The CMG on dsDNA is blocked by two non-homologous duplex arms that form the T-DNA structure (**see Fig. 1a**). Our recent report showed that CMG melts the two non-homologous arms, demonstrating that CMG tracks on dsDNA with force (Langston and O’Donnell, 2019). DNA tracking enzymes interact with the charged phosphate backbone, and to determine if CMG binds only one strand while tracking on dsDNA, our earlier study inserted 10 nucleotides having methyl groups on the phosphodiester backbone (methylphosphonate), a strategy previously used to demonstrate that a phage DNA packaging motor tracks on only one strand while encircling dsDNA (Aathavan et al., 2009). The previous results with CMG using a 10mer section of neutral DNA on one strand or the other showed that CMG interacts 2-3 times more favorably with the 3’-5’ tracking strand compared to the non-tracking 5’-3’ strand. We tried 30 neutral linkages, but this appeared to stop CMG activity regardless of the strand they were on. In the current report we increase the length of neutral DNA on the T-DNA from 10 to a 20mer tract, about the length of the central channel of CMG. The result shows an approximate 8-to 10-fold difference between unwinding the T-DNA with a 20mer neutral segment on the tracking compared to the non-tracking strand (**Fig. 1**). The use of DNA lacking methylphosphonate linkages is shown for comparison; unwinding is nearly the same as T-DNA with a 20mer neutral DNA on the non-tracking strand. These results confirm our earlier results, but greatly strengthen the conclusion that CMG tracks on only one strand while encircling dsDNA.

**Figure 1.**
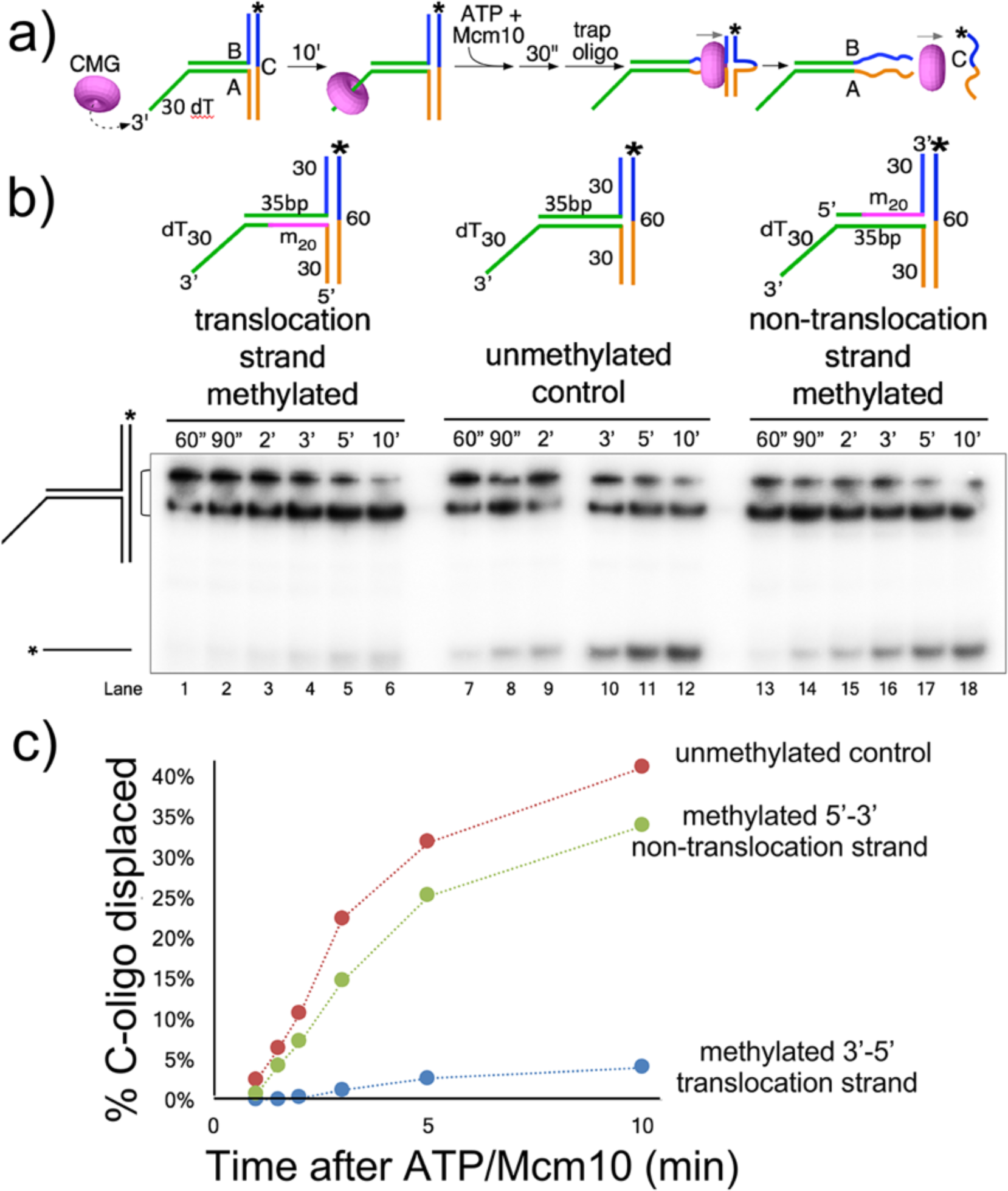
CMG tracks on one strand while encircling double-strand DNA. **a)** The reaction scheme. CMG has been shown to load onto T-DNAs via a 3’ ssDNA dT tail and melt the cross-bar oligo C off the non-homologous arms of Oligos A/B (Langston and O’Donnell, 2019). Mcm10 is also present in these reactions, but is not illustrated for clarity. **b)** Either oligo A or B contains a 20mer section of neutral DNA having uncharged methylphosponate linkages. A time course of unwinding is shown in the gel. C) Quantitation shows that unwinding is severely inhibited when the 3’-5’ tracking strand (Oligo A) contains the neutral DNA (blue circles), but when the non-tracking strand (Oligo B) contains the neutral DNA (green circles), unwinding is nearly the same as T-DNA lacking neutral DNA on either strand (red circles). Values are the average of three independent experiments and the error bars show the standard deviation.

### Only two CMGs can bind the “origin mimic” DNA using AMPPNP

To load head-to-head CMGs onto dsDNA we placed two 3’ ssDNA dT tails on both ends of a 150 bp duplex, which we refer to here as “origin mimic” DNA (**Fig. 2**). The origin mimic DNA was used in our earlier study to demonstrate DNA melting by head-to-head CMGs in an Mcm10 dependent reaction, while control DNAs having only one 3’ tail on one end or the other were not unwound (Langston and O’Donnell, 2019). Since ATP is needed for CMG to load onto DNA, the earlier report incubated ATP with CMG and DNA for only 45 s, to limit DNA CMG loading, before adding a ssDNA trap to shut down further CMG loading. However, we could not be sure whether more than 2 CMG+Mcm10 loaded in 45 s. Earlier studies with Drosophila and human CMG showed that ATPγs or AMPPNP can be substituted for ATP to support DNA binding in EMSA assays (Ilves et al., 2010; Kang et al., 2012). We recently found that ATP*γ*S supports *S. cerevisiae* CMG unwinding of a 30mer duplex fork (Yuan et al., 2019), and therefore decided to examine reactions using AMPPNP.

**Figure 2.**
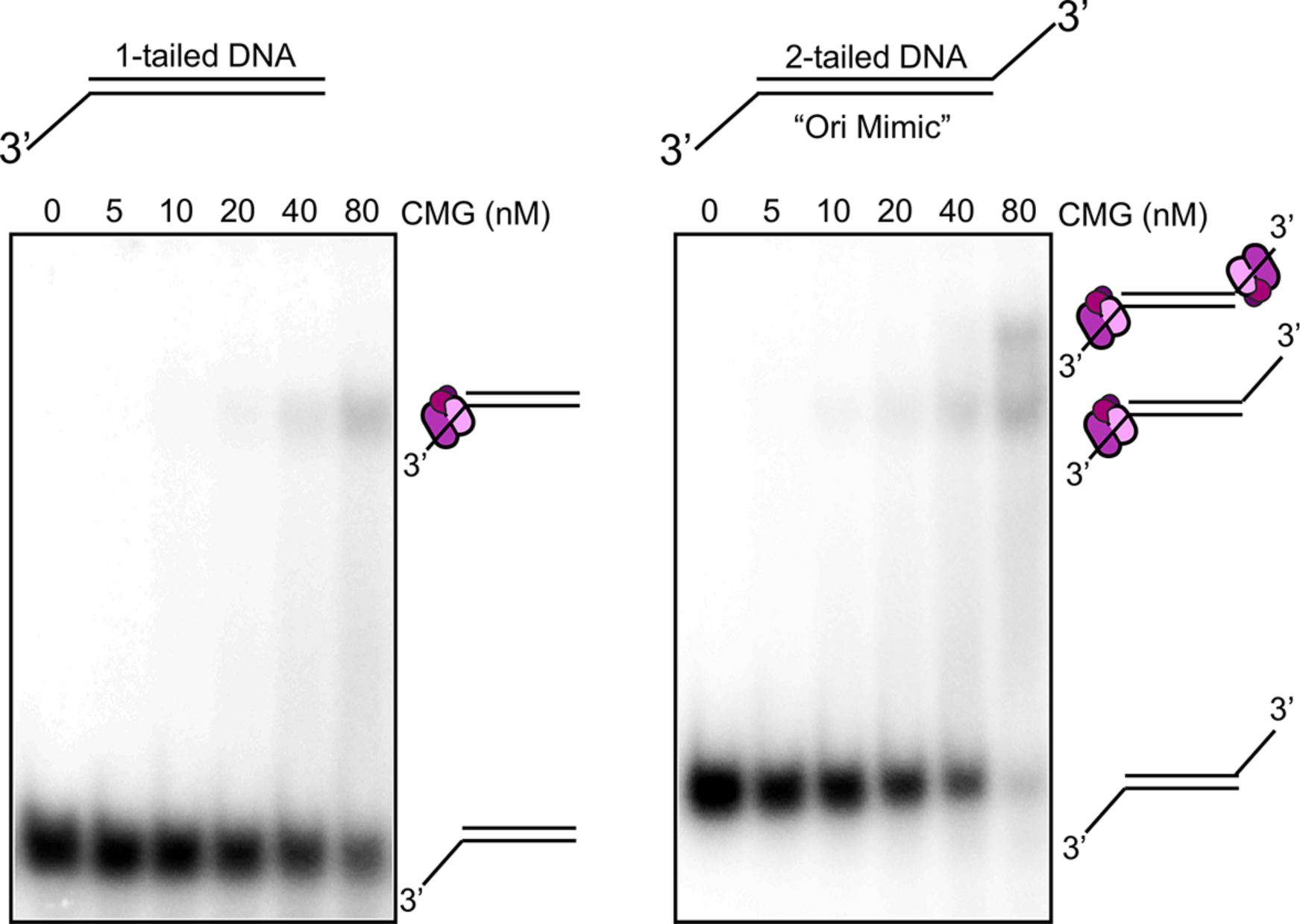
Only 2 CMGs can bind “Ori Mimic” DNA using AMPPNP. EMSA assays were performed using AMPPNP for loading CMG onto 150 bp duplexes containing either one or two 3’ single-strand dT tails. CMG was titrated into reactions containing DNA and 0.2 mM AMPPNP, then analysed for gel shifts by neutral PAGE. Left: The DNA containing only one 3’ tail gave only one gel shift band. Right: The DNA containing two 3’ tails gave two gel shift bands. The presumed interpretation of the bands is shown to the right of the gels. EMSA assay reactions were repeated twice with the same result.

We earlier observed by cryoEM that AMPPNP enables CMG to thread onto a 3’ end, and suggested CMG could not track beyond the 3’ end without hydrolysable nucleotide (Georgescu et al., 2017). This predicts that AMPPNP will only allow 2 CMGs to bind the origin mimic, one on each end. To test this, we performed EMSA binding assays using CMG and AMPPNP with a goal to determine the maximum stoichiometry of CMG binding to the origin mimic two 3’ tailed DNA. We also compared it to a single 3’ tailed DNA of the same sequence, but having only one 3’ tail. Mcm10 was omitted from this experiment because it binds both ssDNA and dsDNA (Warren et al., 2008) and causes DNA shifts. The results show that the origin mimic with two tails can bind either one, or two CMG, but no more (**Fig. 2**). The same DNA, but having only one 3’ tail, gave only one EMSA gel shift band (**Fig. 2**). The results indicate that the maximum occupancy of CMG on the origin mimic with AMPPNP is 2 CMG.

### DNA unwinding by two head-to-head CMGs that encircle dsDNA is efficient

Having observed that non-hydrolysable AMPPNP provides a maximum binding of two CMGs to the 2 tails of the origin mimic substrate enabled us to perform the unwinding assay with a longer preincubation than the short 45 s with ATP in our earlier report. Hence, we gave a 10 minute pre-incubation of CMG, Mcm10, origin mimic DNA and 0.2 mM AMPPNP, then initiated the reaction by adding a solution containing 5 mM ATP and a ssDNA trap that prevents further CMG loading in the presence of ATP. The results, in **Fig. 3**, show a more vigorous unwinding reaction than our earlier study (Langston and O’Donnell, 2019), likely due to enabling more time for two CMGs to bind the two opposing 3’ ssDNA tails of the origin mimic DNA. The trap prevents unwinding when added before proteins (**Fig. 3**), shown previously (Langston and O’Donnell, 2019).

**Figure 3.**
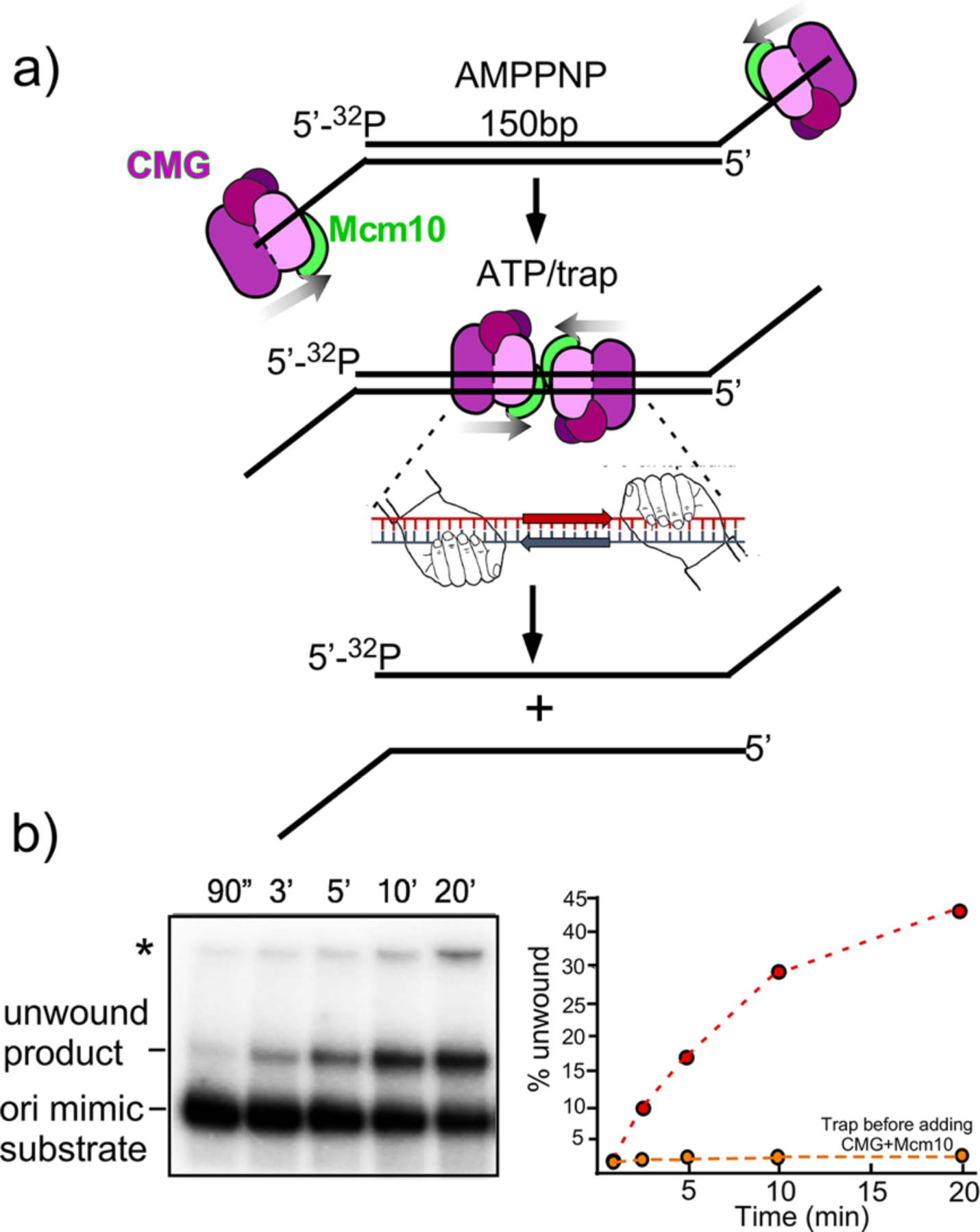
Preloading CMG-Mcm10 on DNA with AMPPNP gives robust unwinding of the origin mimic DNA. **a)** Scheme of the reaction. CMG+Mcm10 are preincubated with AMPPNP and an origin mimic DNA containing two 3’ ssDNA tails. DNA unwinding was initiated upon adding a mixture of ATP and inhibitory trap ssDNA. The two hands in the inset of the middle diagram indicate that the 2 CMGs bind, and thus apply force, to opposite strands of dsDNA. **b)** Gel analysis of a time course of unwinding, three independent trials are shown. The unwound DNA and origin mimic are identified to the left of the gel. Controls from our previous study showed that both 3’ tails are required to observe unwinding (Langston and O’Donnell, 2019). The asterisk indicates a gel-shift of the substrate by CMG and Mcm10. **C)** Quantitation of the % unwinding at each time point in the gel (red dashed line), the error bars represent one standard deviation of the triplicate data. The orange dashed line is the result of adding trap DNA before proteins (data from Figure supplement 1 for Figure 4 in (Langston and O’Donnell, 2019)).

## DISCUSSION

We have shown earlier that dsDNA is unwound by head-to-head CMG motors that encircle dsDNA, likely by shearing the DNA since each opposing CMG binds one strand (i.e. its tracking strand) tighter than the other strand (Langston and O’Donnell, 2019). Single-molecule studies have shown that pulling opposite ends of dsDNA to 65 pN results in dsDNA melting to form ssDNA (King et al., 2016; van Mameren et al., 2009). There are at least two precedents for motor oligomers that encircle dsDNA and track on it with nearly 65 pN, the phi29 phage packaging motor (Smith et al., 2001) and the FtsK chromosome partitioning motor of *E. coli* (Pease et al., 2005). In the case of head-to-head CMGs, there would be two CMGs and thus each would only need contribute half the 65 pN force needed to melt dsDNA.

### Proposed mechanisms of the CMG ds-to-ss transition

In **Fig. 4**, we illustrate steps needed for CMGs to transition from encircling dsDNA to encircling ssDNA, a prerequisite for CMGs to leave the origin and produce bidirectional forks. Shearing dsDNA, shown here and earlier (Langston and O’Donnell, 2019) involves both the breaking of hydrogen bonds between base pairs and the removal of the turns in DNA before CMG can transition to ssDNA. By way of illustration these two actions are illustrated as two separate steps in **Fig. 4**, but they likely occur in a concerted reaction. The two CMGs encircling dsDNA require Mcm10 for ability to move from the origin (**Fig. 4a**,**b**) (reviewed in Bell and Labib, 2016). Upon Mcm10 binding, the turns in DNA are illustrated as being removed (**Fig. 4b**,**c**). The double hexamer of Mcm2-7 encloses about 6 turns (60 bp) of DNA (Abid Ali et al., 2017; Noguchi et al., 2017). As force is applied by the head-to-head CMGs pulling on opposite strands, the 6 turns would likely diffuse out the C-tier ends of the CMGs as overwound DNA for topoisomerases to remove (Postow et al., 1999). By way of illustration, opposite rotations of each CMG are indicated in Fig. 4 to remove the turns, but CMG rotation may not necessarily be required. Of note, CMG is demonstrated to untwist about 0.7 turn in the absence of Mcm10 (Douglas et al., 2018). The second step is disruption of base pairing, illustrated in **Fig. 4d**. These steps would be followed by expulsion of the non-tracking strand from each CMG (**Fig. 4e**), and then passage of the two CMG for departure from the origin and formation of bidirectional forks (**Fig. 4f**).

The motor PS1 loops are nearly at the N-C-domain intersection, and one may question how much DNA must be melted for the CMG ds-ss transition. There are at a minimum 3 important considerations for this event: **1)** CMG-Mcm10 may continue to unwind, producing ssDNA loops that may extrude out the back (C-tier) of the CMGs. 2) If the CMGs let go of the ssDNA, at least transiently, this could provide a sufficient length of ssDNA for both CMGs to expel their respective non-tracking strand (**Fig. 4f**). **2)** CMG may only need to expel a strand through the N-tier ring of Mcm2-7, which is a much more rigid entity than the C-tier (Yuan et al., 2016). The Mcm4/6 interface of the N-tier has the least buried area and has been proposed as a likely position for N-tier ring opening (Yuan et al., 2016). Once one strand is expelled from the N-tier, the C-motors would continue translocating, forming more ssDNA. The ATP driven conformation changes needed to translocate on DNA occur in the C-tier, and these changes may provide ample opportunity for ssDNA passage through C-tier interfaces. **3)** ssDNA, when stretched, is longer than dsDNA, and thus conversion of dsDNA to ssDNA between the PS1 motor loops of the head-to-head CMGs may be sufficient for CMG to transit from ds to ss DNA via a ssDNA gap along the length of CMG. Indeed, a ssDNA gap that enables CMG to transit from ds-to-ssDNA (and from ss-to-ds DNA) has recently been demonstrated (Wasserman et al., 2019).

**Figure 4.**
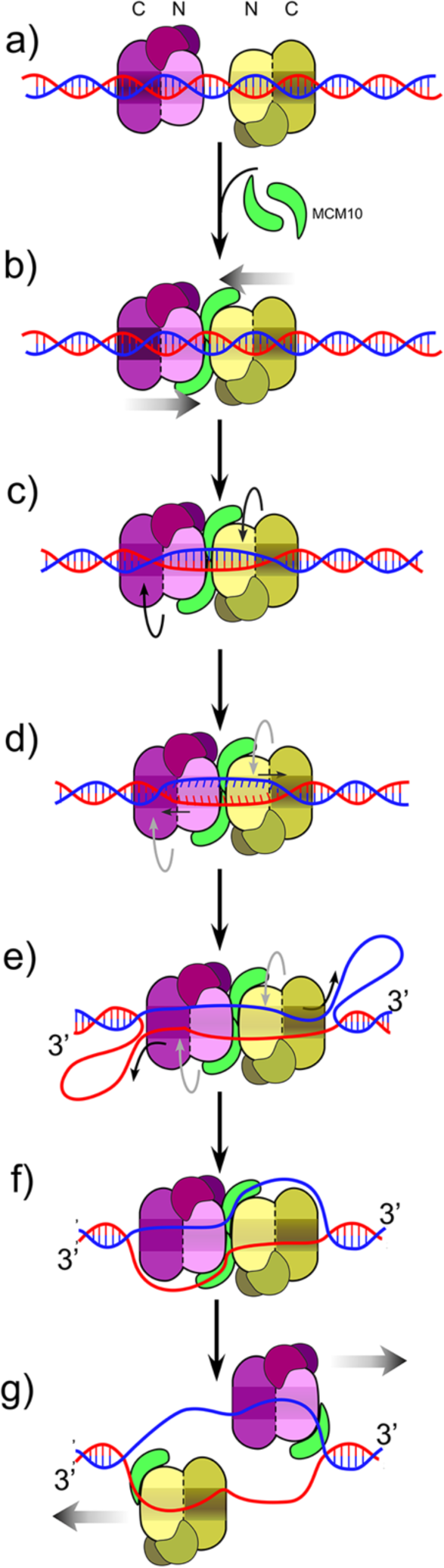
Illustration of DNA melting by head-to-head CMG motors while encircling dsDNA. **a)** Head-to-head CMGs are assembled onto dsDNA at origins, and **b)** Mcm10 binds CMG which is required for CMGs to move from the origin, but the CMG-Mcm10s are directed inward, blocking one another. **c)** The force generated by ATP driven movement of inward directed CMGs causes DNA to move instead. Turns are illustrated to be removed by CMGs that rotate in opposite directions, as a consequence of CMGs using ATP hydrolysis to track toward one another, yet being blocked by one another’s presence. **d)** Base-pairing is removed as a consequence of the same force described for panel c. Note: actions in panels c and d likely occur in a concerted fashion, and are only illustrated separately to show that removal of turns and base pairing both need to occur for CMG to transition from ds-to-ssDNA. **e)** Extensive DNA unwinding by continued DNA shearing in which the motors pull on opposite strands, making each strand move in opposite directions, would form ssDNA loops that exit the C-tier of CMGs. **e)** CMG-Mcm10 has a ssDNA gate that enables a ds-to-ss transition (Wasserman et al., 2019) illustrated here to be used to expel the non-tracking strand from each CMG, enabling the two CMGs to move past one another **(f)** and establish bidirectional replication forks.

### Is DNA shearing used elsewhere in DNA metabolism?

DNA duplex opening by opposed dsDNA motors could possibly be used to generate ssDNA in other cellular processes of DNA metabolism. An example of a replicative function that was found later to generalize to numerous other types of DNA transactions is the case of DNA sliding clamps, originally identified for their ability to provide processivity to replicative DNA polymerases (Kong et al., 1992; Stukenberg et al., 1991). It is now well known that sliding clamps, like PCNA, are used by a large variety of proteins in DNA metabolism (Georgescu et al., 2015). Furthermore, the sliding clamp also represented the initial finding of a protein that functions by topologically encircling DNA. We now know that many proteins encircle DNA for function. Topological binding was predicted to generalize to other proteins in the report that identified the initial discovery of topological binding by a DNA sliding clamp (Stukenberg et al., 1991). However, the identification of proteins that act by DNA encirclement could not be predicted *a priori*. Likewise, with the ability of head-to-head motors that unwind dsDNA. We are unable to predict where this process may be used beyond origin initiation. We do point out however, that there is a “MCM paradox” in which the number of Mcm2-7 complexes associated with chromatin far outnumber origins (Bell and Labib, 2016). Perhaps these Mcm2-7 complexes are used for processes that are distinct from DNA replication. It will be quite interesting to see in the future if the principles of origin unwinding by opposing dsDNA motors generalize to other, as yet unforeseen processes.

## ACKNOWLEDGMENTS

The authors wish to thank Daniel Zhang for purification of CMG and Nina Y. Yao for artwork of Figs. 2-4.

## Author Contributions

L.L. and R.E.G. performed the experiments, L.L. and M.E.O. oversaw the project. M.E.O. wrote the manuscript.

## Data and materials availability

Materials are available upon reasonable request to M.E.O.

## METHODS

### Reagents and Proteins

Radioactive nucleotides were from Perkin Elmer and unlabeled nucleotides were from GE Healthcare. DNA modification enzymes were from New England Biolabs. CMG and Mcm10 were overexpressed and purified as previously described (Georgescu et al., 2014; Langston et al., 2017; Langston et al., 2014). Protein concentrations were determined using the Bio-Rad Bradford Protein stain using BSA as a standard. DNA oligonucleotides were from Integrated DNA Technologies except for those with methylphosphonate linkages which were from Biosynthesis (Lewisville, TX).

### DNA substrates

For all radiolabeled oligonucleotides, 10 pmol of oligonucleotide was labeled at the 5’ terminus with 0.05 mCi [γ-^32^P]-ATP using T4 Polynucleotide Kinase (New England Biolabs). For annealing, 4 pmol of the radiolabeled strand was mixed with 6 pmol of unlabeled complementary strand, NaCl was added to a final concentration of 200 mM, and the mixture was heated to 90°C and then cooled slowly (e.g. 60 min) to 23°C. DNA oligonucleotides used in this study are listed in Table I.

**Table I.**
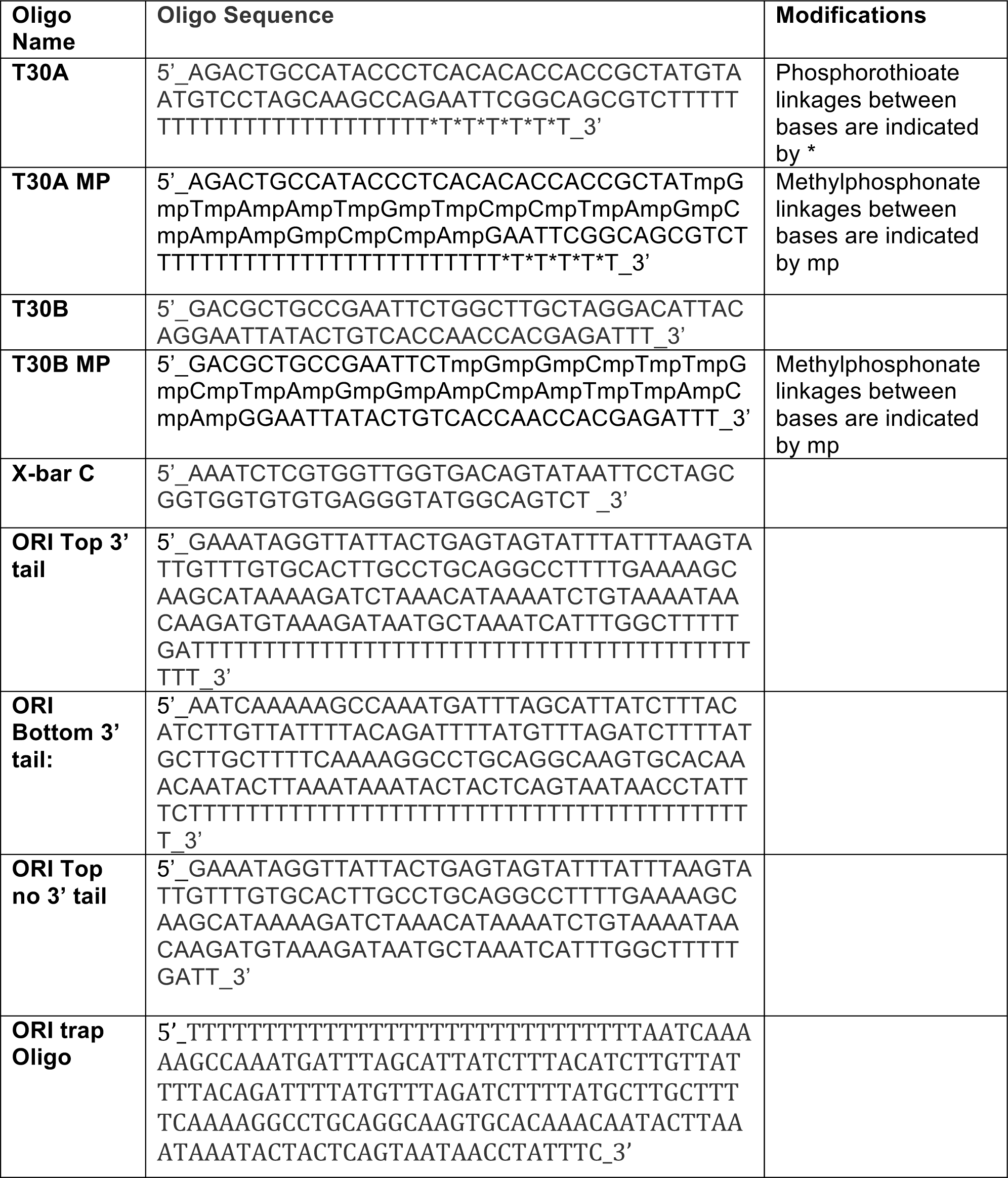
DNA sequences used in this study.

The T-DNA substrate for Fig. 1 was made by annealing unlabeled strands A and B to radiolabeled cross-bar strand C, producing a T-DNA with a 3’ ssDNA of 30 dT, a ds of 35bp, and two non-homologous arms of 30bp each. The same A and B oligonucleotide sequences, but with 20 methylphosphonate linkages in the duplex immediately preceding the T-junction were substituted to make T-DNAs with a neutral section on either the A or the B strand, as illustrated in Fig. 1 (see Table I). For Fig. 3, the origin mimic DNA with two 3’ ssDNA tails was made by annealing unlabeled “ORI Bottom 3’ tail” to radiolabeled “ORI Top 3’ tail”. The origin mimic DNA for EMSA assays was made the same way, except the ORI Bottom 3’ tail oligo was radiolabeled.

### EMSA assays

The ORI Bottom 3’ tail oligo was radiolabeled and used in two separate annealing reactions with either ORI-Top no tail or ORI-Top 3’ tail oligos, along with ORI Bottom 3’ tail oligo, to yield the single-3’ tailed DNA and the origin mimic two 3’-tailed DNA, respectively. Both substrates were subsequently PAGE-purified to eliminate any free contaminating oligo. Binding reactions were performed by incubating 0.5 nM ^32^P-DNA with increasing amounts of CMG (as indicated) in a 10 µL reaction containing 20 mM Tris-acetate, 8% glycerol, 0.02 mM EDTA, 10 mM Na-acetate 10 mM MgSO_4_ and 0.2 mM AMP-PNP. Reactions were incubated 60 min at 30°C, then directly loaded on a 4 % native PAGE gel in TBE buffer containing 5 mM MgSO_4_. Electrophoresis was performed at 4°C at 240 V for 45 min in TBE buffer supplemented with 5 mM MgSO_4_. Gels were wrapped in plastic and exposed to a phosphor screen that was scanned on a Typhoon 9400 laser imager (GE Healthcare).

### T-DNA assays of CMG strand bias

For the T-DNA assays in Fig. 1, reactions contained 20 nM CMG, 40 nM Mcm10, 0.5 nM labeled T-DNA in 20 mM Tris-acetate pH 7.6, 5 mM DTT, 0.1 mM EDTA, 10 mM MgSO_4_, 20 mM KCl, 40 µg/ml BSA and 1 mM ATP. CMG was pre-incubated with the DNA substrate for 10 minutes at 30°C in the absence of ATP in a final volume of 65 µl, and the reaction was started by addition of ATP/Mcm10. To prevent re-annealing of unwound radiolabeled DNA, 20 nM of an unlabeled version of the radiolabeled oligo was added as a trap 30 seconds after starting the reaction. At the indicated times, 10 µl aliquots were removed and stopped with 4 µl of 0.1M EDTA, 5% SDS, 25% glycerol, and 0.01% each of xylene cyanol and bromophenol blue, and flash frozen in liquid nitrogen. Upon completion of the reaction time course, quenched reactions were thawed and loaded on 10% native PAGE minigels by electrophoresis at 100V for 75 minutes in TBE buffer. Gels were washed in distilled water, mounted on Whatman 3MM paper, wrapped in plastic and exposed to a phosphor screen that was scanned on a Typhoon 9400 laser imager (GE Healthcare). Scanned gels were analyzed using ImageQuant TL v2005 software.

### Origin mimic DNA unwinding assays

For study of origin mimic DNA unwinding in Fig. 3, the Ori 3’ top strand was labeled with ^32^P. Reactions containing 40 nM CMG and 80 nM Mcm10 were pre-incubated with 0.5 nM origin mimic DNA, a 150 bp duplex containing two 3’ 30mer dT tails, in 55 µl of 20 mM Tris-acetate pH 7.6, 5 mM DTT, 0.1 mM EDTA, 10 mM MgSO_4_, 20 mM KCl, 40 µg/ml BSA and 0.2 mM AMPPNP for 10 minutes at 30°C. Reactions were initiated upon adding 20 nM unlabeled Ori Trap Oligo (Table I) and 5 mM ATP. The trap oligo binds to the unwound radiolabeled DNA, forming a forked structure that shifts it to a unique position in the native PAGE gel. At the indicated times, 10 µl aliquots were removed and stopped by addition of 4 µl of 150 mM EDTA/7% SDS. 1 µl Proteinase K was added to each quenched reaction and incubated 10’ at 30° C after which 3 µl 0.1M EDTA, 5% SDS, 25% glycerol, and 0.01% each of xylene cyanol and bromophenol blue was added and processed as described above for reactions of Fig. 1 except that electrophoresis was at 100V for 120 minutes.

## Literature Cited

Aathavan, K., Politzer, A.T., Kaplan, A., Moffitt, J.R., Chemla, Y.R., Grimes, S., Jardine, P.J., Anderson, D.L., and Bustamante, C. (2009). Substrate interactions and promiscuity in a viral DNA packaging motor. Nature 461, 669–673.

Abid Ali, F., Douglas, M.E., Locke, J., Pye, V.E., Nans, A., Diffley, J.F.X., and Costa, A. (2017). Cryo-EM structure of a licensed DNA replication origin. Nature Communications 8, 2241.

Bell, S.P., and Labib, K. (2016). Chromosome duplication in Saccharomyces cerevisiae. Genetics 203, 1027–1067.

Bleichert, F., Botchan, M.R., and Berger, J.M. (2017). Mechanisms for initiating cellular DNA replication. Science 355, eaah6317.

Douglas, M.E., Ali, F.A., Costa, A., and Diffley, J.F.X. (2018). The mechanism of eukaryotic CMG helicase activation. Nature 555, 265–268.

Evrin, C., Clarke, P., Zech, J., Lurz, R., Sun, J., Uhle, S., Li, H., Stillman, B., and Speck, C. (2009). A double-hexameric MCM2-7 complex is loaded onto origin DNA during licensing of eukaryotic DNA replication. Proc Natl Acad Sci U S A 106, 20240–20245.

Georgescu, R., Yuan, Z., Bai, L., de Luna Almeida Santos, R., Sun, J., Zhang, D., Yurieva, O., Li, H., and O’Donnell, M.E. (2017). Structure of eukaryotic CMG helicase at a replication fork and implications to replisome architecture and origin initiation. Proceedings of the National Academy of Sciences 114, E697–E706.

Georgescu, R.E., Langston, L.D., O’Donnell, M. (2015). A proposal: Evolution of PCNA’s role as a marker of newly replicated DNA. DNA Repair, 29, 4–15.

Goswami P, Abid Ali F, Douglas ME, Locke J, Purkiss A, Janska A, Eickhoff P, Early A, Nans A, Cheung AMC, Diffley JFX, Costa A. (2018) Structure of DNA-CMG-Pol epsilon elucidates the roles of the non-catalytic polymerase modules in the eukaryotic replisome. Nat Commun. 9:5061.

Ilves, I., Petojevic, T., Pesavento, J.J., and Botchan, M.R. (2010). Activation of the MCM2-7 helicase by association with Cdc45 and GINS proteins. Mol Cell 37, 247–258.

Kang, Y.H., Galal, W.C., Farina, A., Tappin, I., and Hurwitz, J. (2012). Properties of the human Cdc45/Mcm2-7/GINS helicase complex and its action with DNA polymerase epsilon in rolling circle DNA synthesis. Proc Natl Acad Sci U S A 109, 6042–6047.

King, G.A., Peterman, E.J.G., and Wuite, G.J.L. (2016). Unravelling the structural plasticity of stretched DNA under torsional constraint. Nature Communications 7, 11810.

Kong, X.P., Onrust, R., O’Donnell, M., and Kuriyan, J. (1992). Three dimensional structure of the beta subunit of E. coli DNA polymerase III holoenzyme: a sliding DNA clamp. Cell, 69, 425–437.

Langston, L.D., and O’Donnell, M.E. (2017a). Action of CMG with strand-specific DNA blocks supports an internal unwinding mode for the eukaryotic replicative helicase. eLife 6, e23449.

Langston and O’Donnell (2019) An explanation for origin unwinding in eukaryotes. eLife 9:e46515

Lyubimov, A.Y., Strycharska, M., and Berger, J.M. (2011). The nuts and bolts of ring-translocase structure and mechanism. Current Opinion in Structural Biology 21, 240–248.

Meagher, M., Epling, L.B., and Enemark, E.J. (2019). DNA translocation mechanism of the MCM complex and implications for replication initiation. Nat Commun 10, 3117.

Moyer, S.E., Lewis, P.W., and Botchan, M.R. (2006). Isolation of the Cdc45/Mcm2-7/GINS (CMG) complex, a candidate for the eukaryotic DNA replication fork helicase. Proc Natl Acad Sci U S A 103, 10236–10241.

Noguchi, Y., Yuan, Z., Bai, L., Schneider, S., Zhao, G., Stillman, B., Speck, C., and Li, H. (2017). Cryo-EM structure of Mcm2-7 double hexamer on DNA suggests a lagging-strand DNA extrusion model. Proceedings of the National Academy of Sciences. 114, E9529–E9538

O’Donnell, M., Langston, L., and Stillman, B. (2013). Principles and concepts of DNA replication in bacteria, archaea, and eukarya. Cold Spring Harb Perspect Biol 5.

O’Donnell, M.E., and Li, H. (2018). The ring-shaped hexameric helicases that function at DNA replication forks. Nature Structural & Molecular Biology 25, 122–130.

Parker, M.W., Botchan, M.R., and Berger, J.M. (2017). Mechanisms and regulation of DNA replication initiation in eukaryotes. Critical Reviews in Biochemistry and Molecular Biology 52, 107–144.

Pease, P.J., Levy, O., Cost, G.J., Gore, J., Ptacin, J.L., Sherratt, D., Bustamante, C., and Cozzarelli, N.R. (2005). Sequence-Directed DNA Translocation by Purified FtsK. Science 307, 586–590.

Postow, L., Peter, B.J., and Cozzarelli, N.R. (1999). Knot what we thought before: the twisted story of replication. BioEssays 21, 805–808.

Remus, D., Beuron, F., Tolun, G., Griffith, J.D., Morris, E.P., and Diffley, J.F. (2009). Concerted loading of Mcm2-7 double hexamers around DNA during DNA replication origin licensing. Cell 139, 719–730.

Smith, D.E., Tans, S.J., Smith, S.B., Grimes, S., Anderson, D.L., and Bustamante, C. (2001). The bacteriophage φ29 portal motor can package DNA against a large internal force. Nature 413, 748–752.

Stukenberg, P.T., Studwell-Vaughan, P.S., O’Donnell, M. (1991). Mechanism of the sliding beta-clamp of DNA polymerase III holoenzyme. J Biol Chem, 266, 11328–11334.

van Mameren, J., Gross, P., Farge, G., Hooijman, P., Modesti, M., Falkenberg, M., Wuite, G.J.L., and Peterman, E.J.G. (2009). Unraveling the structure of DNA during overstretching by using multicolor, single-molecule fluorescence imaging. Proceedings of the National Academy of Sciences 106, 18231–18236.

Warren EM, Vaithiyalingam S, Haworth J, Greer B, Bielinsky AK, Chazin WJ, Eichman BF. (2008) Structural basis for DNA binding by replication initiator Mcm10. Structure. 16, 1892–901.

Wasserman, M., Schauer, G.D., O’Donnell, M.E. and Liu, S. (2019) Replisome preservation by a single-stranded DNA gate in the CMG helicase. bioRxiv 368472.

Yuan, Z., Bai, L., Sun, J., Georgescu, R.E., Liu, J., O’Donnell, M.E., and Li, H. (2016). Structure of the eukaryotic replicative CMG helicase suggests a pumpjack motion for translocation. Nat Struct Mol Biol, 23, 217–24.

Yuan, Z., Georgescu, R., de Luna Almeida, S.R., Zhang, D., Bai, L., Yao, N.Y., Zhao, G., O’Donnell, M.E., and Li, H. (2019) Ctf4 organizes sister replisomes and Pol *α* into a replication factory. BioRxiv 735746.

